# Metabolism-driven, high-efficiency mining of ethanol-tolerant microorganisms from pit mud microbiota using Raman flow cytometry

**DOI:** 10.64898/2026.02.13.705682

**Authors:** Teng Xu, Qing Sun, Gongchao Jing, Yongming Duan, Xiaohang Wang, Xinyun Yi, Changle Wu, Huizi Zhu, Jian Xu, Xiaowei Zheng, Xixian Wang, Jia Zhang

## Abstract

The discovery of stress-tolerant microorganisms from complex microbiomes is frequently constrained by low screening throughput and the inability of culture-based approaches to access single-cell functional phenotypes, particularly for rare but highly resilient taxa. Here, we establish a metabolism-driven *screen-before-culture* strategy by integrating D_2_O-labelled single-cell Raman spectroscopy (SCRS) with Raman-activated cell sorting (RACS) to directly target ethanol-tolerant cells based on metabolic vitality. By quantifying carbon-deuterium (C-D) incorporation as a single-cell readout of de novo anabolic activity under ethanol stress, this platform enabled culture-independent enrichment of a rare, high-vitality subpopulation (∼0.2% abundance) from pit mud microbiomes at a sorting throughput of ∼2,400 cells per hour. Using this approach, six highly ethanol-tolerant strains were successfully isolated, whereas parallel conventional culture-first screening of the same samples yielded predominantly low-tolerance isolates, with an overall screening efficiency of only 22.22%. Rapid single-cell ethanol tolerance assessment based on SCRS, completed within 7 h, showed that all sorted strains exhibited strong tolerance to 8% (v/v) ethanol, with Raman Tolerance Index (RTI) values exceeding 50%. Among them, *Lactiplantibacillus plantarum* F4 displayed the highest tolerance (RTI = 85.05 ± 3.41%). Comparative transcriptomic analyses of representative strains revealed mechanistically coherent ethanol adaptation strategies, including ethanol-derived carbon recycling, dynamic membrane lipid remodeling, and reinforced redox homeostasis. These responses directly underpin the metabolic activity captured by the Raman screening signal, validating its physiological relevance. This integrated SCRS-RACS workflow achieved orders-of-magnitude higher screening throughput, a 4.5-fold improvement in sorting accuracy, and a 6.86-fold increase in assessment efficiency compared with conventional methods. This study establishes a versatile, metabolism-based paradigm for the targeted mining of rare, stress-tolerant microorganisms from complex microbiomes, with broad implications for industrial biotechnology and microbial ecology.

## Background

Microorganisms function as efficient cellular factories, capable of transforming low-cost renewable feedstocks (including lignocellulose, crude glycerol, and bioethanol) into high-value bioproducts, thereby demonstrating considerable potential for sustainable biomanufacturing [1,2]. According to data from the Organization for Economic Co-operation and Development (OECD), microbial manufacturing already constitutes approximately 40% of the global bioeconomy [3]. Furthermore, propelled by advances in synthetic biology, microbial cell factories are projected to supply 30% of the global demand for chemicals, biofuels, and materials from renewable resources by 2030 [4]. Among diverse microbially synthesized products, bioethanol stands out as a representative clean energy fuel, with its production critically reliant on microbial fermentation [5–7]. However, ethanol accumulation during fermentation imposes severe physiological stress on microbial hosts, leading to alterations in membrane composition, structure, and function, changes in cell morphology, inhibition of cell division, decreased viability, and reduced metabolic activity [8,9]. Consequently, ethanol stress represents a primary bottleneck limiting the commercial feasibility and process intensification of microbial biofuel production [10].

To develop robust microbial chassis with enhanced ethanol tolerance, resilient natural ecosystems offer valuable sources of evolutionary solutions. The pit mud used in traditional Chinese Baijiu fermentation represents a unique natural bioreactor that has been maintained under prolonged high-ethanol and anaerobic conditions for decades. Microorganisms inhabiting this environment have undergone sustained selective pressure, evolving specialized adaptive mechanisms that make pit mud microbiomes an exceptional reservoir for mining ethanol-tolerant strains [11,12].

Despite its potential, the complex structure and high species diversity of pit mud microbiomes present a significant obstacle to the efficient discovery of high-performance strains. Three core technical challenges currently impede progress: (*i*) *Low efficiency of mining*: traditional *culture-first* methods (e.g., dilution-to-extinction in 96- or 384-well plates) are labor-intensive and low-throughput. This approach predominantly isolates dominant taxa, often failing to capture rare, highly ethanol-tolerant phenotypes, which results in poor overall screening efficiency [13,14]. (*ii*) *Slow assessment speed*: ethanol-tolerant strains are typically screened using conventional methods such as optical density (OD) measurement and colony counting, which rely on culture and are time-consuming [15,16]. (*iii*) *Insufficient evaluation depth*: growth-based assays yield delayed, population-averaged data. Such methods are ineffective at detecting cells that enter a viable but non-culturable (VBNC) state under stress and lack the resolution to accurately distinguish strain-specific phenotypic differences in response to ethanol challenge [17,18].

To overcome these limitations, D_2_O-based single-cell Raman spectroscopy (SCRS) provides a powerful strategy to identify and assess the ethanol-tolerant cells by measuring the single-cell metabolic vitality under ethanol stress. Metabolically active cells incorporate deuterium from D_2_O into newly synthesized biomolecules, generating a detectable carbon-deuterium (C-D) signature in their SCRS. The intensity of this C-D peak serves as a quantitative indicator of cellular metabolic vitality, indicating the ethanol tolerance when cultured under ethanol conditions [18–20]. When coupled with Raman-activated cell sorting (RACS), SCRS enables label-free, non-invasive, and broadly applicable high-throughput screening of functionally superior microbial cells [21–23]. Previous RACS platforms, including positive dielectrophoresis-induced deterministic lateral displacement-based RACS (pDEP-DLD-RACS) [24–26] and flow-mode optical tweezers-based RACS [27,28], have successfully profiled and isolated rare functional cells from random mutagenesis libraries and complex gut microbiota, highlighting the feasibility of applying this strategy to pit mud microbiomes.

Here, we established an integrated high-throughput ethanol-tolerant cells screening workflow by coupling D_2_O-SCRS with RACS. SCRS was first used to quantify metabolic vitality within pit mud microbiota under ethanol stress, enabling the identification of highly active, ethanol-tolerant cells at single-cell resolution. These target cells were subsequently enriched by RACS, followed by cultivation and validation. Using this metabolism-driven *screen-before-culture* strategy, six ethanol-tolerant strains were successfully isolated, all of which exhibited strong tolerance to 8% (v/v) ethanol, with Raman Tolerance Index (RTI) values exceeding 50%. Notably, two low-abundance strains, F4 and F5, displayed exceptional ethanol tolerance supported by extensive transcriptomic remodeling, particularly involving ethanol-derived energy conservation and membrane lipid biosynthesis. Together, these results demonstrate that integrating D_2_O-SCRS with RACS provides a generally applicable platform for mining stress-tolerant microorganisms from complex microbiomes and offers a powerful route for engineering robust microbial chassis for synthetic biology and industrial biotechnology.

## Materials and methods

### Chemicals, Strains, and Sample Collection

MRS medium was purchased from Hope Bio-technology Co., Ltd. (Qingdao, China). PBS (1×, pH 7.4, 0.01 M) was obtained from Solarbio Science & Technology Co., Ltd. (Beijing, China). Tween 20 was sourced from Sangon Biotech Co., Ltd. (Shanghai, China). Sodium pyrophosphate was purchased from Aladdin Biochemical Technology Co., Ltd. (Shanghai, China). Ethanol was supplied by Sinopharm Chemical Reagent Co., Ltd. (Shanghai, China). Deuterium oxide (D_2_O) for cell labelling was acquired from Sigma-Aldrich (USA).

The strains CICC 6009, BNCC 194390, *E. coli* 35218, and the DH5α-GFP strain were preserved in the laboratory culture collection. All strains were stored in 20% (v/v) glycerol at -80 ℃. For routine cultivation, *E. coli* 35218 and DH5α-GFP were grown in LB medium at 37 °C with shaking at 180 rpm. Strains CICC 6009 and BNCC 194390 were cultured in MRS medium at 37 °C under facultative anaerobic conditions. All cultures were harvested at the mid-logarithmic growth phase for subsequent experiments.

The pit mud samples were provided by COFCO Corporation and collected from fermentation pit. A systematic sampling strategy was implemented to comprehensively represent the microbial community. Samples were collected from the central bottom, four corners, as well as the middle and top sections of the pit walls. All samples were divided into two groups according to depth: the surface layer (A-F, 0-5 cm) and the deep layer (G-L, 5-15 cm), with 10 g collected per sample. After collection, samples were immediately aliquoted into sterile centrifuge tubes, supplemented with glycerol at a final concentration of 20% (v/v) as a cryoprotectant, and stored at -20 ℃ for subsequent microbial isolation and cultivation.

### Microbial Cell Extraction and Purification from Pit Mud

To obtain a purified microbial cell solution suitable for direct Raman-activated cell sorting (RACS), cells were extracted from pit mud samples using a Nycodenz density gradient centrifugation protocol, which effectively separates microbial cells from particulate matter. Initially, samples of composite pit mud from different locations across the fermentation pit were fully homogenized. A 3 g portion of the homogenized sample was suspended in 15 mL of sterile PBS with 0.5% (v/v) Tween 20 and agitated at room temperature for 30 minutes to disperse cells. The slurry was centrifuged at a low speed (50 ×*g*, 5 min) to remove big soil particles. The primary cell suspension’s supernatant was gathered, The leftover pellet was re-extracted with 10 mL of PBS with 0.5% (v/v) Tween 20 by stirring for one minute, followed by a second low-speed centrifugation step at 50 ×*g* for 5 minutes, and the supernatant was combined with the primary suspension.

The combined crude cell extract was filtered through a sterile 40 µm nylon cell strainer to remove remaining coarse debris. To further purify, 6 mL of the filtrate was carefully layered over an equal volume of 80% (w/v) Nycodenz solution (in triplicate) and centrifuged at 14,000 ×*g* for 1 hour at 4 ℃. After centrifugation, the distinct cell band at the sample-Nycodenz interface was carefully aspirated. An equal volume of ddH_2_O was added to this aspirate to dilute the Nycodenz medium, and cells were pelleted by a final centrifugation step (10,000 ×*g*, 5 min, 4 ℃). To avoid cross-contamination, all equipments were rigorously washed with ddH_2_O and 75% ethanol between each sample.

### Conventional plate-counting assay for ethanol tolerance

For the plate-counting assay, strains BNCC 194390 and CICC 6009 were assessed with and without ethanol stress (15%, v/v). Unstressed controls were plated directly after serial dilution to obtain baseline colony counts (*N_0h_*). Stressed samples were treated with 15% (v/v) ethanol in MRS medium for 24 hours before plating, yielding post-stress counts (*N_24h_*). Three biological replicates were performed per condition. The survival rate (SR) for each strain was calculated as:

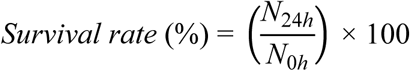

### Development of a Raman-based ethanol tolerance assessment method using D_2_O labelling

To validate the SCRS-based ethanol tolerance assessment method, cultures of the two strains (BNCC 194390 and CICC 6009) from the same batches used in the conventional plate-counting assay were processed in parallel. The SCRS-based workflow comprised three main steps (**Fig. 2A**): sample preparation, automated SCRS acquisition, and data analysis.

Step (*i*): Sample preparation. Cultures were standardized to an OD_600_ of 1.0 and divided into three treatment groups for stress and D_2_O-labelling incubation. All groups requiring D_2_O incubation were subjected to a uniform concentration of 75% D_2_O with an incubation volume of 100 μL per group. Specifically, Group 1 had no ethanol treatment and no D_2_O incubation. Group 2 received no ethanol treatment and was incubated in D_2_O for 6 hours. Group 3 was treated with ethanol (15%, v/v) for 24 hours, followed by incubation in 75% D_2_O for 6 hours.

Step (*ii*): Automated SCRS acquisition. For each sample, the Raman acquisition was conducted using a 532 nm laser power of 80-100 mW, and at least 200 SCRS were collected.

Step (*iii*): Data analysis. Raman spectral data processing and quantitative analysis were performed in R software (v4.1.3). Preprocessing steps included baseline correction and smoothing were applied to Raman spectra. The CDR was calculated based on the preprocessed spectra, which was derived by dividing the C-D peak area (2040-2300 cm^-1^) by the sum of the C-D and C-H peak areas (2800-3100 cm^-1^). This CDR value was then used to calculate the RTI for each strain according to the formula:

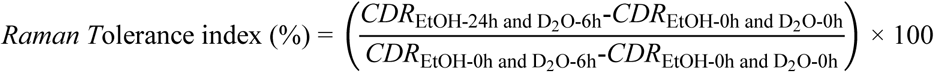

### System setup and validation of RACS

The RACS chip was fabricated using quartz by bonding a bottom thick layer and a top thin cover slice (200 μm in thickness) with double-sided tape (50 μm in thickness). (**Fig. 3B**). The structure of the double-sided adhesive tape was designed using AutoCAD 2024 (Autodesk, USA). Cells were loaded via a PEEK tube with a small inner diameter (≈305 μm; Cole-Parmer, USA), which was connected to the microfluidic device and an air pump. Raman microscopy was performed on a FlowRACS instrument (Qingdao Single-Cell Biotechnology Co., Ltd., CN). The FlowRACS instrument employed an Nd:YAG 532 nm laser emitter as the excitation light source, an Nd:YAG 1064 nm laser emitter as the optical tweezers source, a 50× objective (NA = 0.7, Sunny Group CO., LTD., CN) to focus the laser beam on the sample, an electron-multiplying charge-coupled device (EMCCD; Newton DU970N-BV, Andor, UK) to collect SCRS. During the sorting experiments, the loading rates were set to 7 μL/min for the sheath buffer and 6 μL/min for the sample; the Raman acquisition was conducted using a 532 nm laser power of 50 mW, and the acquisition time of 0.5 s.

To assess the accuracy in sorting D_2_O-labelled cells, the *E. coli* 35218 cells without D_2_O incubation (i.e., without CD peak) and the DH5α-GFP cells with D_2_O incubation (i.e., with CD peak) were 9:1 mixed and underwent RACS to sort the DH5α-GFP cells with the sorting criteria of “CDR > 0.03” under 0.5 s acquisition time and 50 mW 532 nm laser. Sorted cells were collected and plated, and the purity of DH5α-GFP cells were counted via fluorescence. From the enumeration results, the sorting accuracy was calculated as *N*_fluorescence_ / (*N*_fluorescence_ + *N_non-_*_fluorescence_).

### Transcriptomic Analysis

For transcriptomic analysis, strains F4 (*Lactiplantibacillus plantarum*) and F5 (*Staphylococcus epidermidis*) were cultured under ethanol stress conditions. Cultures were divided into ethanol-treated (8%, v/v) and control (culture medium without ethanol) groups, with samples collected at 24 h and 48 h. Each condition was prepared with three independent biological replicates. Cells were incubated anaerobically at 37 °C, harvested by centrifugation (5,000 ×*g*, 10 min, 4 °C), immediately frozen in liquid nitrogen. Transcriptomic sequencing was performed by OE Biotech Co., Ltd. (Shanghai, China).

Strand-specific cDNA libraries were constructed and sequenced on the DNBSEQ-T7 platform (MGI, China) using a paired-end strategy. Raw reads were filtered and aligned to the reference genomes of *L. plantarum* WCFS1 and *S. epidermidis* ATCC 12228. Transcriptomic bioinformatic analyses were performed using the BMKCloud platform (https://www.biocloud.net). Differentially expressed genes (DEGs) were identified using DESeq2, with screening criteria of |log₂ fold change| ≥ 1.5 and a false discovery rate (FDR) ≤ 0.05. Principal component analysis (PCA) was conducted to assess transcriptional variation among samples. Functional enrichment analyses of DEGs were performed based on KEGG pathways using a hypergeometric test.

## Results

### Comparison of SCRS-based and conventional screening-evaluation workflows

Conventional isolation of ethanol-tolerant microorganisms from complex microbiomes typically follows a *culture-first-screen-next* workflow, which suffers from major limitations in efficiency, throughput, and phenotypic resolution. Environmental samples are first spread onto solid media and incubated for initial growth (∼48 h), followed by multiple rounds of colony purification to obtain pure isolates, requiring approximately 144-192 h to establish a strain library. Subsequent ethanol tolerance evaluation using growth-based assays, such as plate counting or optical density measurements, adds an additional ∼48 h. Consequently, the entire workflow spans 192-240 h. Moreover, reliance on manual colony picking restricts screening throughput to only hundreds of clones per researcher per day (**Fig. 1A**).

**Fig. 1.**
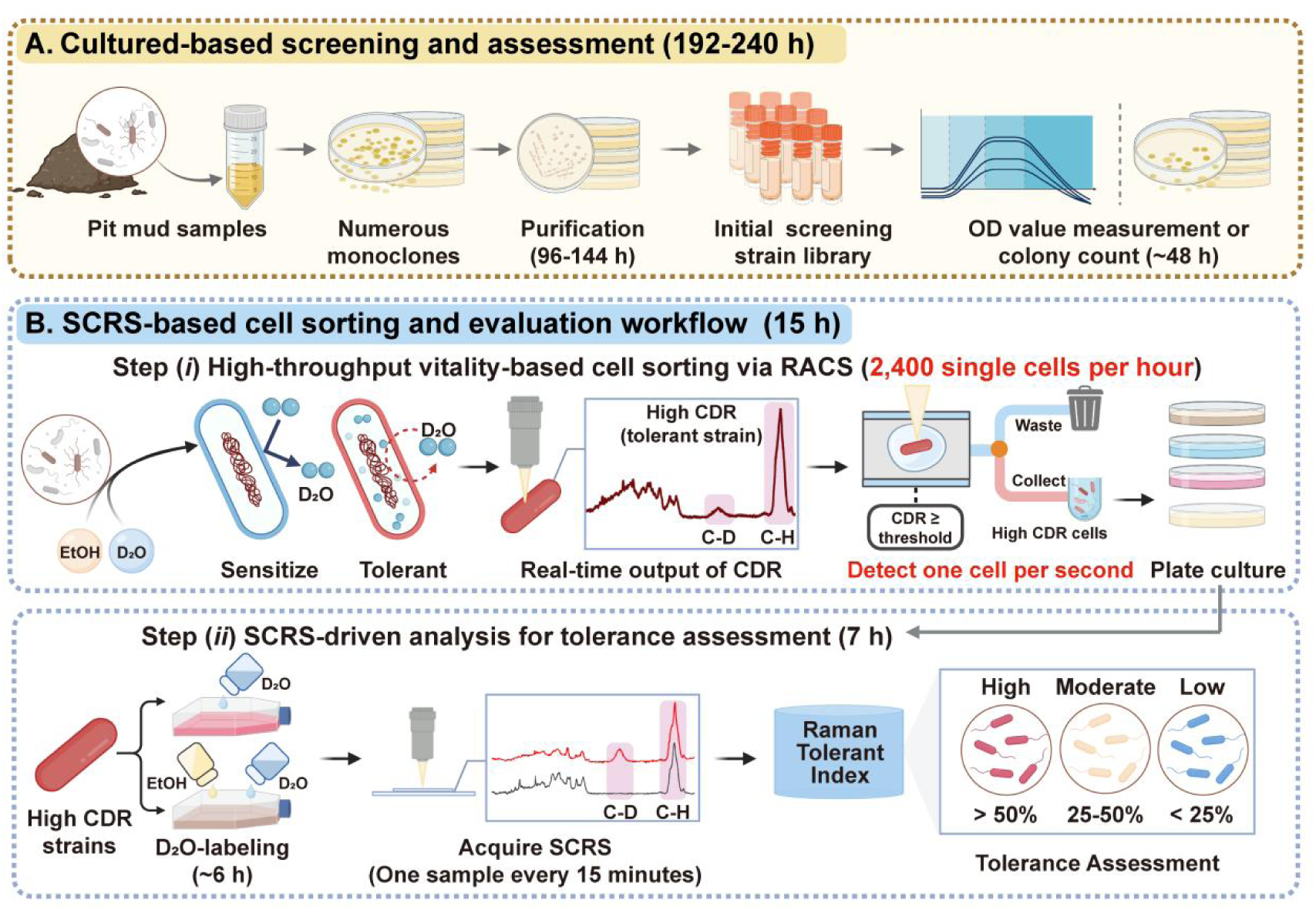
Comparison of conventional *culture-first* and SCRS-based *screen-before-culture* workflows. (A) Conventional *culture-first* screening pipeline for isolating ethanol-tolerant microorganisms from complex microbiomes. This workflow relies on prolonged cultivation, iterative colony purification, and growth-based tolerance evaluation, resulting in low throughput and extended processing time. (B) Metabolism-driven SCRS-based *screen-before-culture* workflow. Single-cell metabolic vitality under ethanol stress is first quantified by D_2_O-labeled single-cell Raman spectroscopy (SCRS), enabling rapid identification and enrichment of rare ethanol-tolerant cells via Raman-activated cell sorting (RACS), followed by targeted cultivation and rapid SCRS-based tolerance evaluation.

To overcome these limitations, we developed a high-throughput, metabolism-driven *screen-before-culture* workflow based on SCRS. This workflow leverages the metabolic vitality of cells under stress, quantified by the intensity of the C-D band after D_2_O labelling, as a sorting criterion. Utilizing a RACS system, microbial communities are scanned at single-cell resolution with a throughput of up to 2,400 cells per hour. By applying a defined C-D ratio (CDR, defined as the ratio between the CD band area (2040-2300 cm^-1^) and the CD plus CH bands (2800-3050 cm^-1^) area) threshold, the system rapidly identifies and enriches rare, high-vitality cells directly from the microbiome, narrowing down thousands of candidates to approximately dozens for cultivation. This targeted sorting and enrichment process is completed within approximately 8 hours (**Fig. 1B; *Step i***). Following cultivation, ethanol tolerance is rapidly evaluated using the same SCRS technology, replacing slow, population-averaged conventional assays. The assessment, which includes a 6-hour D_2_O incubation under stress, automated SCRS acquisition (15 minutes), and data analysis, provides results in under 7 hours. This method yields two key quantitative metrics: metabolic vitality, indicating the strain’s functional resilience, and the Raman Tolerance Index (RTI), which categorizes tolerance levels with single-cell resolution (**Fig. 1B; *Step ii***).

Overall, the integrated SCRS-based workflow completes the entire screening and evaluation cycle within ∼15 h, representing a >15-fold reduction in screening time compared with conventional culture-first approaches (∼8 h vs. ∼144-192 h) and a hundreds-fold increase in screening throughput (∼2,400 cells/h vs. ∼100 clones/day). In addition, the evaluation phase is nearly seven times faster (7 h vs. 48 h) and provides unprecedented quantitative insight into functional heterogeneity at the single-cell level. Together, this efficient and systematic pipeline enables precise discovery and in-depth characterization of ethanol-tolerant microorganisms from complex microbial communities.

### Development and validation of a SCRS-based method for rapid ethanol tolerance evaluation

To address the critical bottleneck of slow and low-resolution tolerance assessment in conventional screening pipelines, we developed a rapid, single-cell phenotypic method based on D_2_O-labelled SCRS. This approach quantifies a strain’s metabolic resilience by measuring its ability to incorporate deuterium into new biomolecules under ethanol stress, providing a direct readout of cellular vitality. The standardized procedure consists of three key steps (**Fig. 2A**): *Step 1. Stress incubation and D_2_O labelling:* cells are subjected to ethanol stress (8% ethanol) or control condition (without ethanol) in a medium containing 75% D_2_O for 6 hours. *Step 2. Automated SCRS acquisition*: an automated system acquires high-quality Raman spectra from individual cells, ensuring high-throughput data collection. *Step 3. Quantitative data analysis*: the CDR values are used to compute the Raman Tolerance Index (RTI), a metric that compares the metabolic activity under stress to non-stress conditions, with a higher RTI indicating greater ethanol tolerance.

**Fig. 2.**
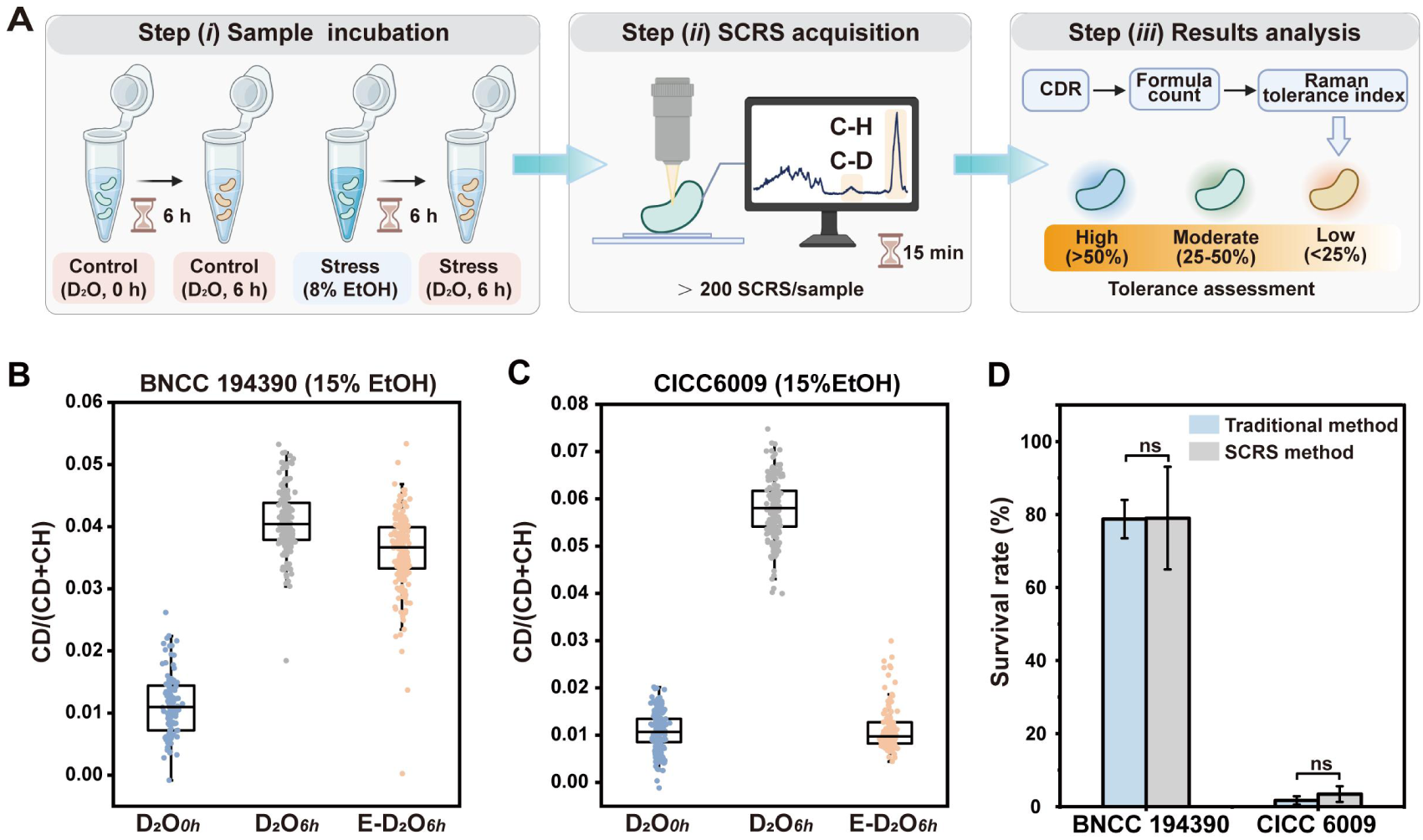
Validation of the SCRS-based ethanol tolerance assessment method. (**A**) Schematic workflow of the D_2_O-labelled SCRS method, comprising three main steps: (*i*) sample preparation, involving culture standardization, ethanol stress treatment, and D_2_O labelling; (*ii*) automated SCRS acquisition; and (*iii*) data analysis for calculating the Carbon-Deuterium ratio (CDR) and Raman Tolerance Index (RTI). The CDR distribution of individual cells from BNCC 194390 (**B**) and CICC 6009 (**C**) strains under different conditions, demonstrating distinct metabolic vitality responses to ethanol stress. Group 1 is the control group (without ethanol stress and D_2_O incubation), abbreviated as D_2_O*_0h_*. Group 2 is the D_2_O-labelled group under non-stress conditions (6 h incubation), abbreviated as D_2_O*_6h_*. Group 3 is the D_2_O-labelled group under ethanol stress (24 h stress followed by 6 h D_2_O incubation), abbreviated as E-D_2_O*_6h_*. (**D**) Bar charts comparing the tolerance levels (RTI or survival rate) of the two strains as quantified by the SCRS method and the traditional plate counting method, respectively. No significant difference (ns, *p* > 0.05) was observed between the two methods, confirming the accuracy of the SCRS-based assessment.

We rigorously validated the accuracy of this SCRS-based RTI against the traditional plate-based colony counting (**Fig. S1**). Using an ethanol-tolerant strain (BNCC 194390) and an ethanol-sensitive strain (CICC 6009) as benchmarks, we observed a clear phenotypic distinction. Under ethanol stress and D_2_O incubation, the CDR distribution of the tolerant BNCC 194390 shifted markedly upward, indicating robust metabolic activity. In stark contrast, the CDR distribution of the sensitive CICC 6009 remained unchanged, revealing severe metabolic inhibition (**Fig. 2B**).

Critically, the quantitative RTI values derived from SCRS showed no statistically significant difference from those obtained via the traditional method (BNCC 194390: 78.72% vs. 79.01%; CICC 6009: 3.43% vs. 1.69%; *p* > 0.05; **Fig. 2B** and **Table S1**). This strong correlation validates the SCRS-based method as a highly accurate alternative, indicating that the SCRS-based assessment method can reduce the evaluation time from several days to ∼7 hours while maintaining the accuracy. Consequently, we established the RTI as our primary tolerance metric and adopted the classification system from Techaparin et al. (2017) [29], categorizing strains as high tolerance (> 50%), moderate tolerance (25%∼50%), or low tolerance (< 25%).

### Design of the RACS system

Previously, a flow-mode optical tweezers-based RACS was developed and realized the sorting of D_2_O-labelled cells from gut microbiota, yet the throughput (3.3-8.3 cells/min) still can be increased further [27,28]. Here, also developed a RACS by integrating optical tweezers with a microfluidic sorter, realizing the sorting of D_2_O-labelled cells with a throughput of ∼40 cells/min. The RACS consists of a self-developed confocal Raman microscope for acquiring the SCRS and a microfluidic chip connected to pump to operate the cell flow (**Fig. 3A)**. To avoid the interference of Raman background from the chip, reduce the cost, and simplify fabrication, the microfluidic chip was assembled by bonding two quartz slices with patterned double-sided tape (**Fig. 3B**) [30]. With this design, only one sheath flow was needed to focus the sample stream horizontally (rather than another sheath flow to focus vertically), which confines the cell flow by default into the waste outlet with a flow ratio of 1.2 : 1 (sheath buffer : sample) (**Fig. 3C**). Moreover, by designing a flow resistance module (i.e., the winding channel) at both waste and collection channel, the stability of the fluid between the waste and collection outlets can be maintained during sorting (**Fig. 3C** and **Video S1**). After loading the cells via inlet 1 and sheath buffer via inlet 2 (**Fig. 3C*i***), the single cells were trapped by optical tweezers produced by 1064 nm laser (**Fig. 3C*ii***), and then moved to Point 1 for acquiring SCRS via 532 nm laser (**Fig. 3C*iii***). The nontarget cells were released directly and the optical tweezers moved back to the cell flow to trap the cells again (**Fig. 3C*iv***). The target cells were further moved and released at Point 2 (**Fig. 3C*v***). To ensure that cells flow fluently into wasted outlet when released at Point 1 and flow fluently into collection outlet when released at Point 2, the angle that separates the waste and collection channel was to align to the middle of the vertical sheath flow channel (**Fig. 3D**).

**Fig. 3.**
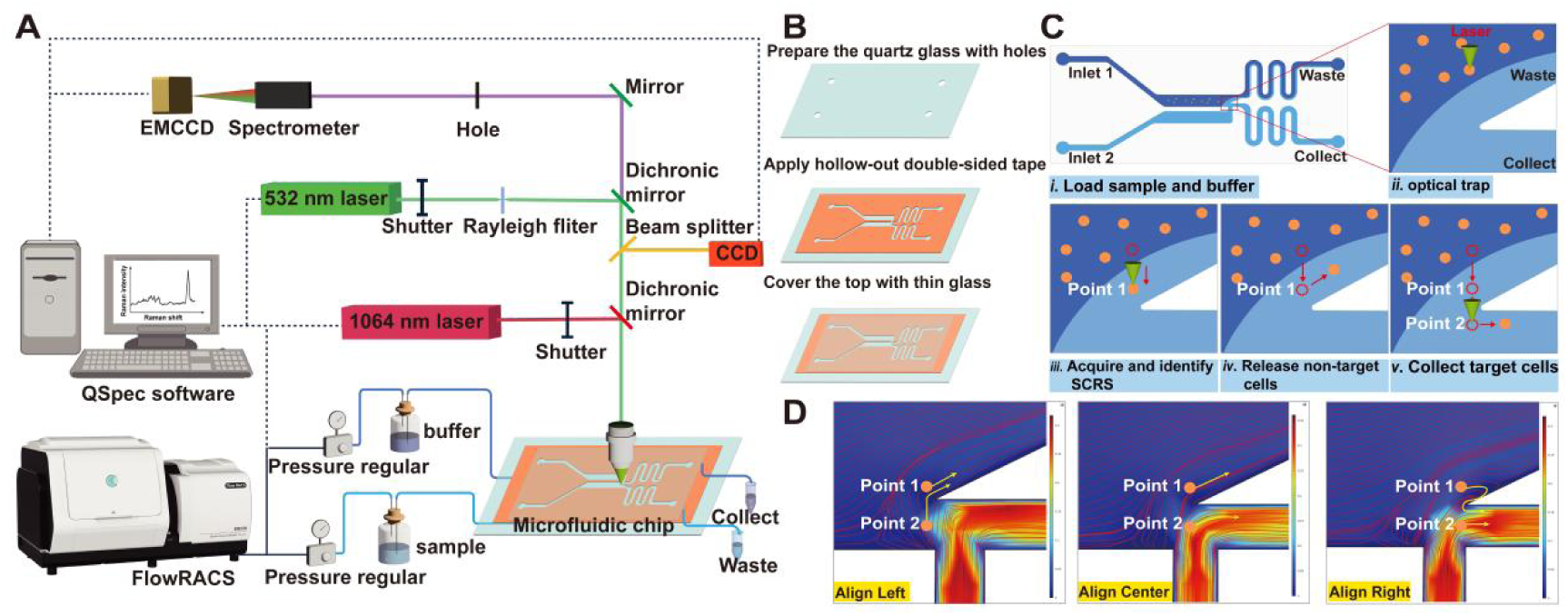
Design and operation of RACS System. (**A**) Schematic of RACS System Setup. (**B**) Schematic of RACS chip design and fabrication. (**C**) System operation. *i.* The sample and buffer were loaded into the channel. *ii.* Single cells were trapped by optical tweezers. *iii.* The trapped single cell was moved to Point 1 for SCRS acquisition and identification. *iv.* The non-target cells were released at Point 1. *v*. The target cells were further moved and released at Point 2. (**D**) Simulate the influence of angle position on fluid motion using COMSOL. Align Left, the cells released at Point 1 flow toward waste outlet, while the cells released at Point 2 flow toward waste outlet. Align Center, the cells released at Point 1 flow toward waste outlet, while the cells released at Point 2 flow toward collect outlet. Align Right, the cells released at Point 1 flow toward collect outlet, while the cells released at Release Point 2 flow toward collect outlet.

### Evaluating performance in sorting D_2_O-labelled cells

To assess the capability of optical tweezers-based RACS in sorting D_2_O-labelled cells, we employed *E. coli* 35218 (without fluorescence) and DH5α-GFP (with fluorescence) as models. Here, the use of DH5α-GFP allowed direct validation of the sorting accuracy by fluorescence. SCRS reveal a prominent C-D peak from 2040 to 2300 cm^-1^ when incubated with 75 % D_2_O, in both *E. coli* 35218 and DH5α-GFP (**Fig. 4A and B**). Intensity of CDR of >90% *E. coli* 35218 cells and all the DH5α-GFP cells are > 0.03 (n > 100) with D_2_O incubation under 0.5 s acquisition time and 50 mW 532 nm laser power, while that of all the cells are < 0.03 without D_2_O incubation (**Fig. 4C**). Finally, the *E. coli* 35218 cells without D_2_O incubation (i.e., without CD peak) and the DH5α-GFP cells with D_2_O incubation (i.e., with CD peak) were 9 : 1 mixed and underwent RACS to sort the DH5α-GFP cells with the sorting criteria of “CDR > 0.03” under 0.5 s acquisition time and 50 mW 532 nm laser. Sorted cells were collected and plated, and the purity of DH5α-GFP cells were counted via Fluorescence. Due to the high efficiency in trapping single cells (**Video S2**) and identifying the DH5α-GFP cells, the sorting accuracy reached to 91.3% (**Fig. 4D**). Thus, our RACS can sort D_2_O-labelled cells efficiently.

**Fig. 4.**
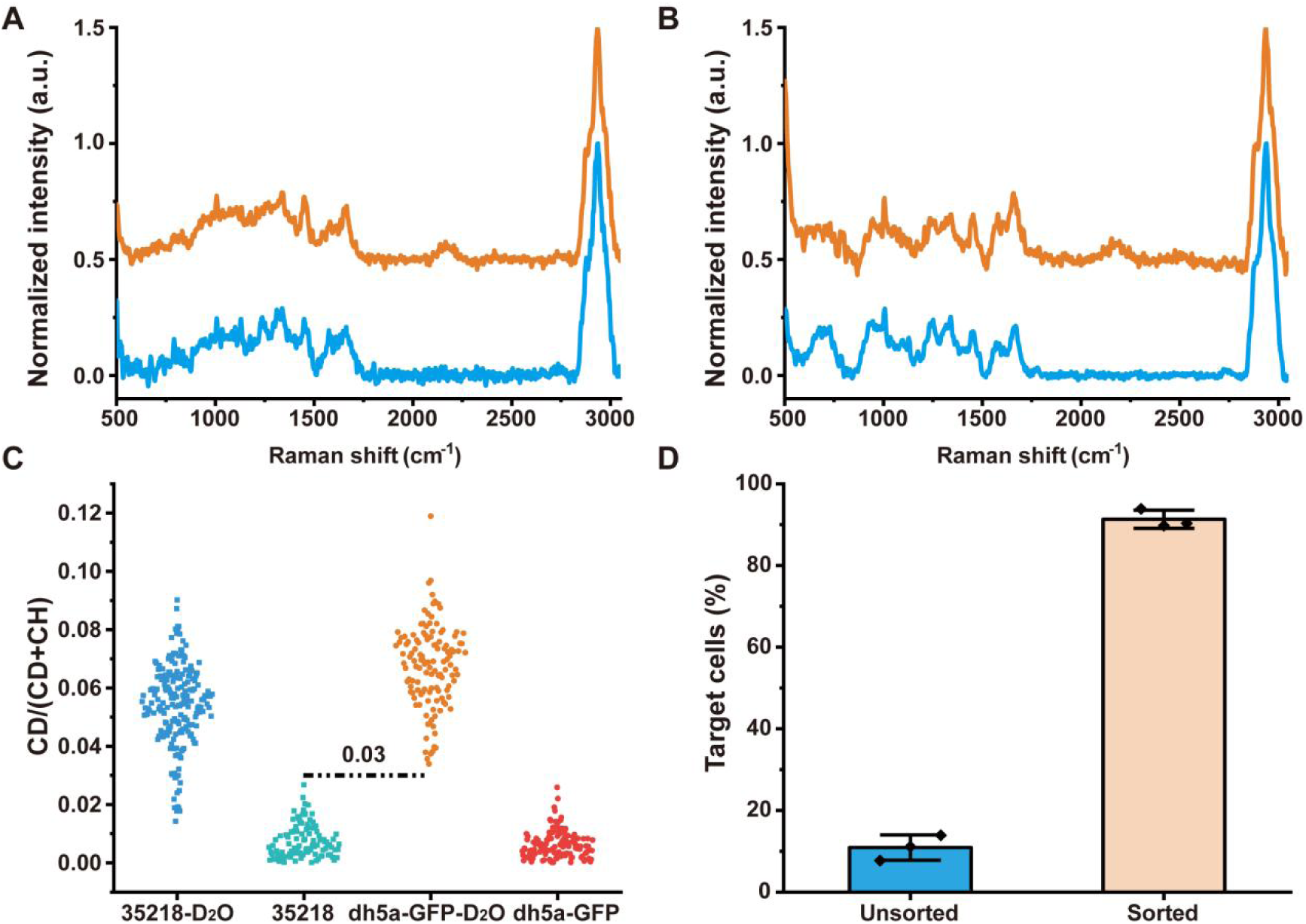
Performance in sorting D_2_O-labelled cells. Average SCRS of *E. coli* 35218 (**A**) and DH5α-GFP (**B**) cells with and without D_2_O treatment. Cells treated with D_2_O exhibit characteristic C-D peaks in the region from 2040 to 2300 cm^-1^, which are absent in non-treated controls. (**C**) Distribution of CDR of *E. coli* 35218 and DH5α-GFP cells with and without D_2_O treatment. The dashed line (CDR = 0.03) indicates the sorting criteria. (**D**) Accuracy in sorting D_2_O-labelled cells. Error bars indicate the SD of three independent experiments.

### Sorting the highly ethanol-tolerant cells from pit mud microbiota

To sort the highly ethanol-tolerant cells from pit mud microbiota, over 1,000 cells were first measured by RACS after D_2_O incubation, while the value of CDR in SCRS that separates the top 1% of cells in metabolic activity under ethanol condition from the remaining 99% of cells was chosen as the sorting criteria (**Fig. S2**). We then applied this RACS-based, *screen-first* strategy in parallel with a conventional *culture-first* approach to the same pit mud sample. The conventional method, relying on random colony picking, yielded nine candidate strains (Y1-Y9). In contrast, the RACS strategy enabled the direct, targeted enrichment of six high-vitality cells (F1-F6) from the pre-identified rare subpopulation. Evaluation of the isolated strains confirmed the efficacy of our targeted approach. Under ethanol stress, all RACS-derived strains (F1-F6) maintained significantly elevated CDR levels compared to their baseline, demonstrating robust metabolic function (**Fig. 5A**). Conversely, most conventionally isolated strains (Y1-Y9) showed no significant increase in metabolic vitality, with only two strains (Y2 and Y5) exhibiting stable performance (**Fig. 5A**).

**Fig. 5.**
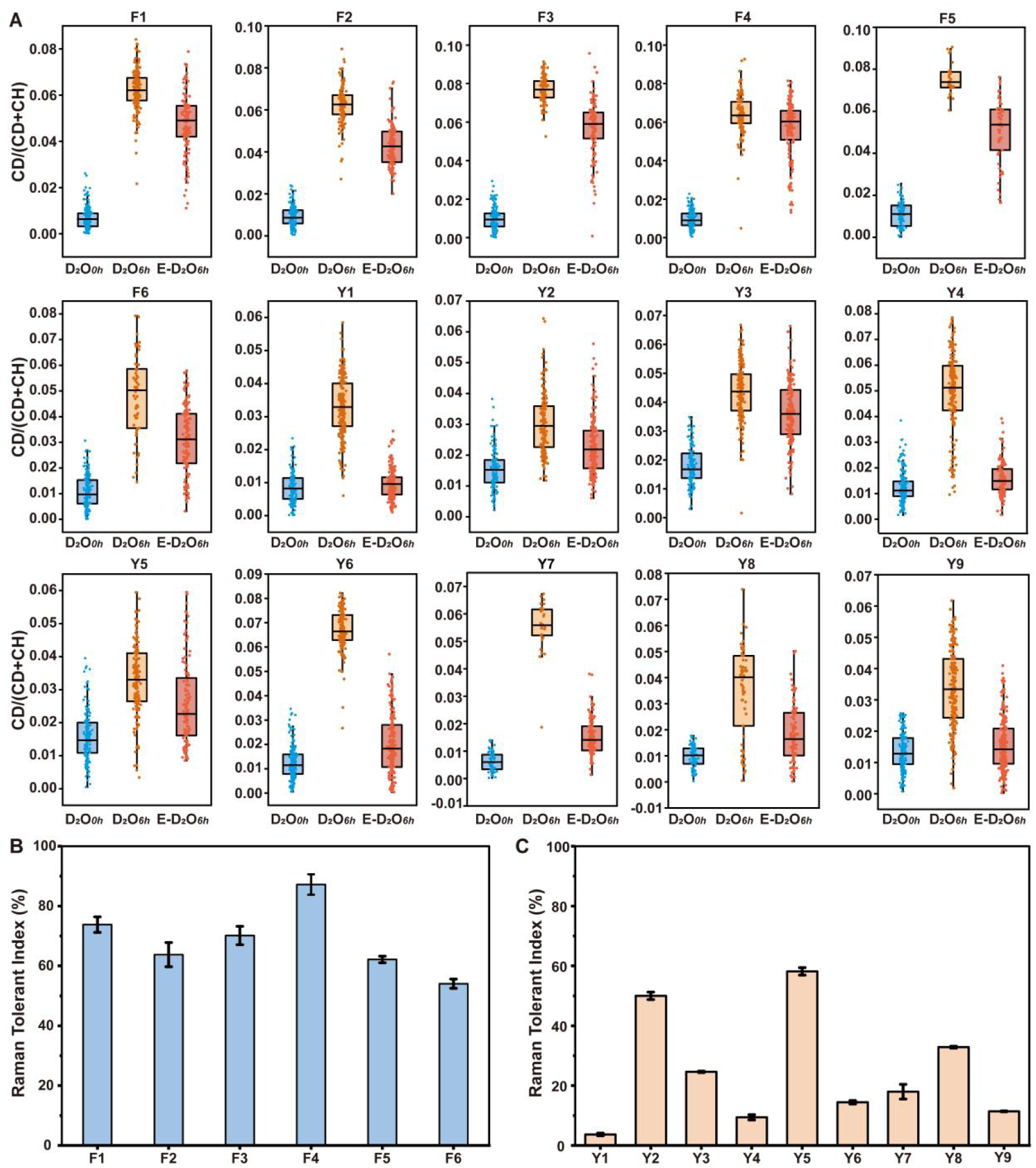
Superior performance of RACS-sorted strains over conventionally screened isolates. (**A**) Metabolic vitality of strains isolated via RACS (F1-F6) and conventional methods (Y1-Y9). Group 1 is the control group (without ethanol stress and D_2_O incubation), abbreviated as D_2_O*_0h_*. Group 2 is the D_2_O-labelled group under non-stress conditions (6 h incubation), abbreviated as D_2_O*_6h_*. Group 3 is the D_2_O-labelled group under ethanol stress (24 h stress followed by 6 h D_2_O incubation), abbreviated as E-D_2_O*_6h_*. Raman tolerance index (RTI) distribution for (**B**) RACS-sorted strain (F1-F6) and (**C**) traditionally isolated strains (Y1-Y9). While most traditional isolates have an RTI < 50%, all RACS-sorted strains have an RTI > 50% (high tolerance).

This performance disparity was quantitatively underscored by the Raman Tolerance Index (RTI; see **Table S2** for complete data). Using RTI ≥ 50% as the criterion for high tolerance, all six RACS-sorted strains were classified as highly tolerant, with *Lactobacillus plantarum* F4 exhibiting the highest RTI of 85.05 ± 3.41% (**Fig. 5B**). In stark contrast, only two out of nine strains from the conventional method met this high-tolerance threshold, resulting in a low screening efficiency of 22.22% (**Fig. 5C**). These results demonstrate that the conventional method is largely stochastic and inefficient for mining rare, high-performance phenotypes, whereas the RACS-based strategy provides a direct and highly efficient route for the targeted discovery of robust ethanol-tolerant strains from complex microbiomes.

### Targeted capture of rare, ethanol-resilient taxa via RACS

A D_2_O-labelled SCRS strategy was employed to selectively enrich highly active cells from pit mud microbiota, using a normalized C-D ratios > 0.08 (top 1% of the population) as the sorting threshold (**Fig. S2**). Using this metabolism-driven criterion, six ethanol-tolerant strains were successfully isolated and subsequently identified by sequencing. All sorted cells were detected in the corresponding bulk pit mud metagenomes, confirming that they originated from the native microbial community rather than culture-derived artifacts.

Quantitative abundance profiling revealed that these strains were present at very low levels in the original community, including F1-F4 (*L. plantarum*, 0.0066%), F5 (*S. epidermidis*, 0.0035%), and F6 (*L. acidipiscis*, 0.19%; **Table S2**). Despite their rarity, all six strains exhibited high ethanol tolerance, indicating that metabolically active but low-abundance taxa can represent functionally important members of pit mud microbiomes, particularly under high-ethanol fermentation conditions. These results highlight the advantage of a metabolism-based, screen-before-culture strategy for uncovering rare yet high-performance microbial phenotypes that are largely inaccessible through conventional random isolation.

### Divergent adaptation of two rare strains under ethanol stress

To elucidate ethanol tolerance strategies in rare taxa, transcriptomic profiling was performed on two low-abundance yet highly ethanol-tolerant strains, F4 (*Lactiplantibacillus plantarum*) and F5 (*Staphylococcus epidermidis*; **Table S2**). Both strains exhibited pronounced transcriptional remodeling in response to ethanol stress, albeit with distinct temporal dynamics. Strain F4 showed a progressive reduction in the number of differentially expressed genes (DEGs) over time, with 1,684 DEGs at 24 h and 1,479 at 48 h (**Table S3; Fig. 6A, B**). In contrast, strain F5 displayed an expanding transcriptional response, with DEG numbers increasing from 1,023 at 24 h to 1,246 at 48 h (**Table S3; Fig. 6C, D**).

**Fig. 6.**
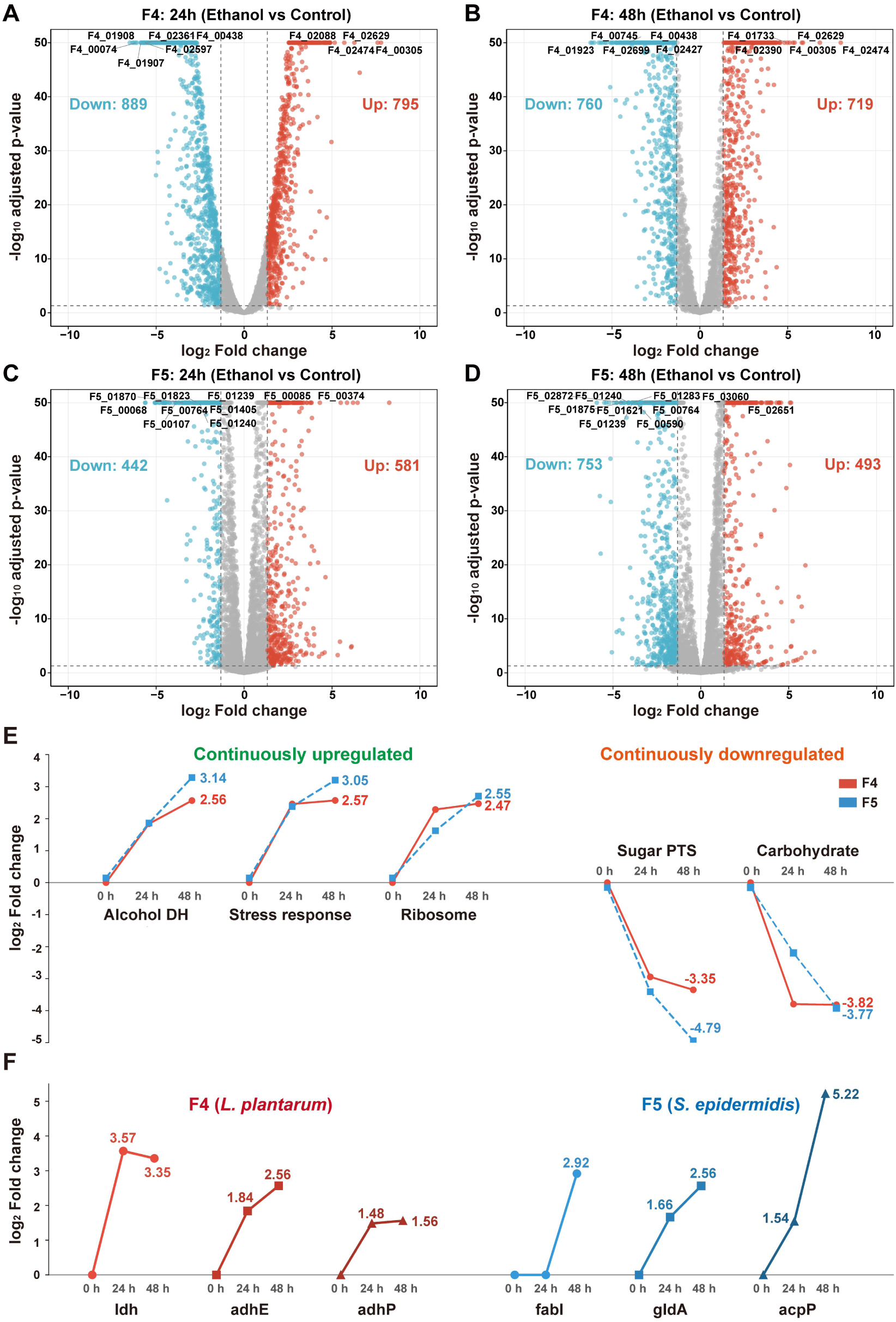
Transcriptional divergence and conservation under ethanol stress. (**A-D**) Volcano plots of differentially expressed genes (DEGs) in *L. plantarum* F4 (**A, B**) and *S. epidermidis* F5 (**C, D**) at 24 h and 48 h. Red and blue dots denote upregulated and downregulated genes, respectively (|log_2_FC| ≥ 1.32), adjusted *p* ≤0.05. (**E**) Conserved monotonic responses. Trends represent the mean log_2_ FC of functional categories consistently upregulated (left) or downregulated (right, e.g., sugar PTS) in both strains. (**F)** Strain-specific adaptive strategies. Expression profiles of key genes driving metabolic sustainment in F4 (fermentation) versus structural reinforcement in F5 (lipid/glycerol metabolism).

Cross-species comparison based on KEGG Orthology (KO) revealed a conserved core ethanol-responsive program. A total of 291 shared KO groups containing DEGs were identified at 24 h, increasing to 300 at 48 h. Among these, 419 gene pairs (47.7%) exhibited concordant regulation at 24 h. Shared responses included the continuous upregulation of alcohol dehydrogenases, stress response, and Ribosome, whereas genes involved in carbohydrate metabolism and sugar phosphotransferase systems (PTS) were progressively downregulated (**Fig. 6E**). Key genes associated with these shared conversed responses are detailed in **Table S4**.

Beyond this conserved core response, the two strains adopted fundamentally different adaptive strategies, reflecting distinct physiological and ecological roles. Strain F5 primarily employed a structural reinforcement strategy, characterized by the monotonic activation of the lipid biosynthesis and glycerol metabolism (*fabI*, *acpP*, *gldA*) to remodel membrane composition to mitigate ethanol-induced osmotic stress (**Fig. 6F**). In contrast, strain F4 prioritized metabolic rerouting to sustain functional activity under ethanol stress. This strategy was marked by strong induction of lactate dehydrogenase (*ldh*), supporting efficient NAD^+^ regeneration and glycolytic flux, while maintaining alcohol dehydrogenases (*adhE*, *adhP*) involved in ethanol catabolism and carbon recycling (**Fig. 6F**).

### Flavor-related metabolic responses of strain F4 to ethanol stress

Building upon this conserved ethanol tolerance framework, we further examined whether strain F4 could maintain functionally relevant metabolic activities under ethanol stress. Notably, beyond stress adaptation, strain F4 retained robust transcriptional activity in pathways associated with flavor precursor formation.

At 24 h, genes involved in organic acid biosynthesis were significantly up-regulated, most prominently *ldh* (encoding L-lactate dehydrogenase), which exhibited a 3.41-fold increase in expression under ethanol exposure. Additionally, aminotransferases, which play key roles in amino acid interconversion and flavor precursor synthesis, exhibited dynamic and selective regulation. Among the 15 aminotransferase genes identified in strain F4, nine were differentially expressed, indicating a selective reshaping of amino acid metabolism. Specifically, gene *araT* (F4 01149) showed the strongest up-regulation (log_2_FC = 4.70 at 24 h, 4.35 at 48 h), whereas gene F4 02788 was significantly down-regulated (log_2_FC = -3.36 at 24 h, -2.17 at 48 h; **Table S5**).

This coordinated transcriptional reprogramming suggests that strain F4 actively prioritizes enzymatic routes supporting organic acid production and amino acid turnover under ethanol stress. Collectively, these findings position strain F4 as a low-abundance yet metabolically highly active member of the pit mud microbiome, capable of sustaining flavor-related metabolic functions even under high ethanol conditions.

## Discussion

The discovery of functionally elite microorganisms from complex communities remains a central challenge in microbiology, particularly when the desired phenotypes are rare, stress-resilient, and uncoupled from rapid growth. Conventional culture-dependent paradigms, reliant on random isolation followed by growth-based screening, are intrinsically inefficient for this task. Reported success rates as low as 3.75% underscore the statistical improbability of capturing rare, high-performance phenotypes through untargeted approaches [34]. This limitation is especially pronounced in structured ecosystems such as pit mud, where our metagenomic analysis revealed that highly ethanol-tolerant strains can constitute less than 1% of the total community. Together, these constraints underscore the need for targeted, function-driven screening strategies.

To address this challenge, we developed a D_2_O-labelled SCRS workflow that fundamentally shifts microbial mining from growth-dependent selection to function-based targeting. The core innovation lies in using metabolic vitality under stress (quantified by the incorporation of deuterium from D_2_O into newly synthesized biomolecules) as a universal and direct screening marker. Unlike strategies that depend on predefined genetic markers or the accumulation of specific metabolites, the C-D Raman signal serves as a direct and generic reporter of *de novo* anabolic activity. This enables unbiased enrichment of functionally superior cells without prior knowledge of genotype, metabolic pathway, or taxonomic identity, substantially expanding the discoverable functional landscape.

The effectiveness of this strategy was demonstrated by the targeted capture of ultra-high-vitality, ethanol-resilient taxa from a complex pit mud microbiome. All sorted strains were confirmed by metagenomic sequencing to originate from the native community, despite their extremely low initial abundance. This highlights a critical advantage of metabolism-driven sorting: it decouples functional importance from numerical dominance, allowing rare but highly active taxa to be systematically recovered. In this context, SCRS-RACS outperforms traditional random isolation not merely in efficiency, but in its ability to directly interrogate functional heterogeneity at the single-cell level.

Multi-omics analyses further provided mechanistic validation for metabolic vitality as a predictive screening criterion. Comparative transcriptomic profiling of two low-abundance yet highly ethanol-tolerant strains, F4 (*Lactiplantibacillus plantarum*) and F5 (*Staphylococcus epidermidis*), revealed a conserved core ethanol-response program alongside distinct adaptive strategies. Shared responses included induction of stress defense pathways, ethanol catabolism, and ribosome biogenesis, accompanied by pronounced repression of phosphotransferase system – mediated sugar uptake and carbohydrate metabolism. These transcriptional features are characteristic of an active, resource-reallocating stress response rather than passive survival.

Beyond this conserved core, the two strains diverged in their dominant ethanol tolerance strategies in a manner that directly explains the mechanistic basis of the SCRS screening signal. Strain F5 primarily adopted a structural reinforcement strategy, characterized by the progressive activation of fatty acid biosynthesis operons and glycerol metabolism genes, supporting membrane remodeling and osmotic protection under ethanol stress. In contrast, strain F4 favored a metabolic sustainment strategy, marked by strong induction of lactate dehydrogenase to promote NAD^+^ regeneration and maintain glycolytic flux, together with sustained expression of alcohol dehydrogenases involved in ethanol catabolism and carbon recycling. Notably, these anabolic and redox-balancing processes are the direct biochemical drivers of deuterium incorporation from D_2_O, indicating that the C-D Raman signal used for cell sorting is not a correlative proxy but a mechanistically grounded spectroscopic readout of the cellular activities that underpin ethanol tolerance.

This tight coupling between single-cell metabolic phenotype and transcriptome-level adaptation establishes a robust scientific foundation for the SCRS-based workflow. By integrating functional screening with molecular interpretation, the approach enables a unified framework linking phenotype, mechanism, and ecological relevance. Notably, the sustained activation of organic acid biosynthesis and amino acid metabolism in strain F4 further indicates that metabolic vitality predicts not only stress resilience but also the capacity to maintain functionally relevant biochemical outputs, such as the production of flavor-related metabolites.

From a practical perspective, the integrated SCRS-RACS platform addresses several long-standing bottlenecks in microbial strain development. The high-throughput sorting capability (∼2,400 cells/hour), combined with culture-free operation, dramatically shortens the discovery timeline. The rapid (<7 h), single-cell resolution tolerance assessment yields quantitative metrics such as the Raman Tolerance Index (RTI), capturing phenotypic heterogeneity that is inaccessible to population-averaged assays based on optical density or colony formation. Together, these features enable a systematic, high-resolution, and mechanism-aware pipeline for mining functional microorganisms.

Nevertheless, the current methodology is best suited for environments containing metabolically active cells. Extending its applicability to ecosystems characterized by widespread dormancy, such as chronically nutrient-limited habitats [36,37], represents an important direction for future development. Potential strategies include the implementation of standardized resuscitation protocols to transiently activate dormant cells prior to sorting [38–41], as well as the integration of advanced computational approaches, such as self-supervised learning, to enhance Raman signal extraction from low-activity cells through improved noise reduction and baseline correction [42,43].

In summary, the D_2_O-labelled SCRS workflow establishes a versatile and mechanism-aware platform that overcomes fundamental limitations of traditional microbiology screening. By providing a direct, culture-independent route from single-cell metabolic phenotype to molecular mechanism, it enables accelerated discovery of robust microbial cell factories. Beyond ethanol tolerance, this approach is readily extensible to mining microorganisms resilient to diverse industrially relevant stresses, including low pH, high salinity, and organic solvents, thereby supporting a paradigm shift toward high-throughput, knowledge-driven exploration of functional microbial resources for sustainable biomanufacturing.

## Conclusion

This study establishes an integrated D_2_O-labelled SCRS platform that fundamentally shifts functional microbial mining from growth-dependent screening to metabolism-driven targeting. By exploiting the carbon-deuterium Raman signal as a direct readout of single-cell metabolic vitality, the platform enabled precise enrichment of a rare yet highly active subpopulation (∼0.2% relative abundance) from a complex pit mud microbiome, leading to the efficient isolation of six strongly ethanol-tolerant strains. Compared with conventional cultivation-based approaches, the SCRS workflow achieved orders-of-magnitude higher screening throughput, a 4.5-fold improvement in sorting accuracy, and a 6.86-fold increase in assessment efficiency.

Importantly, transcriptomic analysis of the top-performing strain, Lactiplantibacillus plantarum F4 (RTI = 85.05 ± 3.41%), demonstrated that the elevated metabolic vitality captured by SCRS is mechanistically underpinned by coordinated stress-adaptive and metabolic reprogramming, thereby validating the physiological relevance of the sorting criterion. Together, these findings establish D_2_O-labelled SCRS as a versatile, culture-independent, and mechanism-aware platform for systematic discovery of high-performance microbial phenotypes. Beyond ethanol tolerance, this strategy is readily extensible to mining microorganisms resilient to diverse industrial stresses, providing a powerful route to accelerate the development of robust microbial cell factories for fermentation, biomanufacturing, and related biotechnological applications.

## Acknowledgments

Not applicable.

## Authors’ contributions

Jia Zhang, Xixian Wang and Xiaowei Zheng designed the research and were involved in writing-reviewing and editing. Teng Xu and Qing Sun performed the experiments and wrote the manuscript. Gongchao Jing analyzed the transcriptomic data. Jian Xu provided critical suggestions. Yongming Duan provided the pit mud samples. Xiaohang Wang, Xinyun Yi, Changle Wu and Huizi Zhu helped with data acquisition. All authors read and approved the final manuscript.

## Ethics approval and consent to participate

Not applicable.

## Consent for publication

Not applicable.

## Data availability statement

The F4 and F5 raw transcriptomic sequencing data have been deposited in the National Center for Biotechnology Information (NCBI) under the BioProject accession number PRJNA1399530 and PRJNA1417456.

