## Supplemental Information for "Metabolism-driven, high-efficiency mining of ethanol-tolerant microorganisms from pit mud microbiota using Raman flow cytometry"

**Supplemental Tables and Figures**

**Table S1. Comparison of SCRS-based and traditional plate counting methods for quantifying strain tolerance (RTI/SR). △CDR: *CDR*_EtOH-24h and D2O-6h_ - CDR_EtOH-0h and D2O-0h_.**

| **Strain** | **Method** | **0 h**  **(Control)** | **24 h**  **(15% ethanol)** | **Tolerance assessment** |
| --- | --- | --- | --- | --- |
| BNCC 194390 | Plate counting (CFU/mL) | 3.02 ± 0.88 × 10^8^ | 2.27 ± 0.20 × 10^8^ | SR: 78.72% ± 5.25 |
|  | SCRS  (△CDR) | 3.17 ± 0.07 × 10^-2^ | 2.49 ± 0.11 × 10^-2^ | RTI: 79.01% ± 14.06 |
|  | *p* value |  |  | H_0_: *p* > 0.05 |
| CICC 6009 | Plat counting (CFU/mL) | 2.27 ± 0.08 × 10^8^ | 8.00 ± 5.35 × 10^6^ | SR: 3.43% ± 2.17 |
|  | SCRS  (△CDR) | 5.08 ± 0.39 × 10^-2^ | 8.99 ± 6.75× 10^-4^ | RTI: 1.69% ± 1.15 |
|  | *p* value |  |  | H_0_: *p* > 0.05 |

**Table S2. Raman tolerance index (RTI) of isolates from RACS-based and conventional methods.**

| **Isolate ID** | **Isolated strains** | **In situ relative abundance (%)** | **Isolation method** | **RTI (%) ± SD** | **Tolerance category** |
| --- | --- | --- | --- | --- | --- |
| F1 | *Lactobacillus plantarum* | 0.0066 | RACS | 73.81 ± 2.61 | High |
| F2 | *Lactobacillus plantarum* | 0.0066 | RACS | 63.78 ± 4.04 | High |
| F3 | *Lactobacillus plantarum* | 0.0066 | RACS | 70.15 ± 3.08 | High |
| F4 | *Lactobacillus plantarum* | 0.0066 | RACS | 85.05 ± 3.41 | High |
| F5 | *Staphylococcus epidermidis* | 0.0035 | RACS | 62.16 ± 1.09 | High |
| F6 | *Ligilactobacillus acidipiscis* | 0.19 | RACS | 54.06 ± 1.54 | High |
| Y1 | / |  | Conventional | 3.68 ± 0.44 | Low |
| Y2 | / |  | Conventional | 50.01 ± 1.25 | High |
| Y3 | / |  | Conventional | 24.61 ± 0.25 | Low |
| Y4 | / |  | Conventional | 9.41 ± 0.87 | Low |
| Y5 | / |  | Conventional | 58.17 ± 1.24 | High |
| Y6 | / |  | Conventional | 14.43 ± 0.58 | Low |
| Y7 | / |  | Conventional | 17.98 ± 2.45 | Low |
| Y8 | / |  | Conventional | 32.86 ± 0.33 | Moderate |
| Y9 | / |  | Conventional | 11.39 ± 0.15 | Low |

**Table S3. Summary of differentially expressed genes (DEGs) under ethanol stress.**

| **Strain** | **Time** | **Up-regulated** | **Down-regulated** | **Total DEGs** |
| --- | --- | --- | --- | --- |
| F1 | 24 h | 795 | 889 | 1,684 |
| F2 | 48 h | 719 | 760 | 1,479 |
| F3 | 24 h | 581 | 442 | 1,023 |
| F4 | 48 h | 493 | 753 | 1,246 |

**Table S4. Shared stress-responsive DEGs in strains F4 and F5.**

| **Category** | **Direction** | **Strain** | **Gene ID** | **Gene_Name** | **log_2_FC**  **(24 h)** | **log_2_FC**  **(48 h)** |
| --- | --- | --- | --- | --- | --- | --- |
| Alcohol dehydrogenase | Up | F5 | F5_03560 | *adh* | 1.71 | 3.14 |
| Alcohol dehydrogenase | Up | F5 | F5_04108 | *gldA* | 1.66 | 2.56 |
| Alcohol dehydrogenase | Up | F4 | F4_03183 | *adhE* | 1.84 | 2.56 |
| Alcohol dehydrogenase | Up | F4 | F4_02644 | *-* | 2.19 | 2.49 |
| Alcohol dehydrogenase | Up | F4 | F4_02734 | *adhP* | 1.48 | 1.56 |
| Ribosome | Up | F5 | F5_01699 | *rpmD* | 1.47 | 2.55 |
| Ribosome | Up | F5 | F5_00825 | *tsf* | 1.79 | 2.26 |
| Ribosome | Up | F5 | F5_00981 | *rbgA* | 1.66 | 2.14 |
| Ribosome | Up | F5 | F5_01717 | *rpsJ* | 1.41 | 1.98 |
| Ribosome | Up | F5 | F5_00824 | *rpsB* | 1.75 | 1.92 |
| Ribosome | Up | F5 | F5_00758 | *rpsT* | 1.76 | 1.78 |
| Ribosome | Up | F4 | F4_01436 | *efp* | 2.29 | 2.47 |
| Ribosome | Up | F4 | F4_01432 | *rplU* | 1.52 | 1.61 |
| Ribosome | Up | F4 | F4_00941 | *rplE* | 1.52 | 1.56 |
| Stress response | Up | F5 | F5_00102 | *usp2* | 2.23 | 3.05 |
| Stress response | Up | F5 | F5_05006 | *nhaX* | 1.81 | 2.45 |
| Stress response | Up | F4 | F4_02002 | *yugI* | 2.46 | 2.57 |
| Stress response | Up | F4 | F4_02082 | *tpx* | 1.6 | 1.63 |
| Carbohydrate metabolism | Down | F5 | F5_01816 | *-* | -1.52 | -1.63 |
| Carbohydrate metabolism | Down | F5 | F5_01483 | *pimB* | -1.67 | -2.21 |
| Carbohydrate metabolism | Down | F5 | F5_01883 | *galM* | -2.05 | -3.77 |
| Carbohydrate metabolism | Down | F4 | F4_02497 | *maa* | -1.67 | -1.74 |
| Carbohydrate metabolism | Down | F4 | F4_02909 | *mutY* | -1.69 | -2 |
| Carbohydrate metabolism | Down | F4 | F4_01777 | *-* | -1.45 | -2.72 |
| Carbohydrate metabolism | Down | F4 | F4_00378 | *lacA* | -2.36 | -2.78 |
| Carbohydrate metabolism | Down | F4 | F4_03126 | *-* | -2.8 | -2.95 |

**Table S4 (Continued from the previous page)**

| **Category** | **Direction** | **Strain** | **Gene ID** | **Gene_Name** | **log_2_FC**  **(24 h)** | **log_2_FC**  **(48 h)** |
| --- | --- | --- | --- | --- | --- | --- |
| Carbohydrate metabolism | Down | F4 | F4_01900 | *-* | -2.94 | -3.34 |
| Carbohydrate metabolism | Down | F4 | F4_03018 | *melA* | -3.79 | -3.82 |
| Carbohydrate metabolism | Down | F4 | F4_03156 | *maa* | -2.72 | -5.32 |
| Carbohydrate metabolism | Down | F4 | F4_03158 | *araD* | -3.91 | -6.01 |
| Carbohydrate metabolism | Down | F4 | F4_03157 | *araA* | -4.04 | -6.13 |
| Sugar PTS | Down | F5 | F5_02777 | *pts31BC* | -1.86 | -2.28 |
| Sugar PTS | Down | F5 | F5_00686 | *srlB* | -1.35 | -2.78 |
| Sugar PTS | Down | F5 | F5_02701 | *pts36C* | -2.84 | -2.98 |
| Sugar PTS | Down | F5 | F5_01950 | *hprK* | -1.69 | -3.35 |
| Sugar PTS | Down | F5 | F5_02703 | *pts36A* | -3.26 | -4.79 |
| Sugar PTS | Down | F4 | F4_00265 | *-* | -1.45 | -1.87 |
| Sugar PTS | Down | F4 | F4_02812 | *fruA* | -2.34 | -2.41 |
| Sugar PTS | Down | F4 | F4_02411 | *pts38A* | -2.69 | -2.91 |
| Sugar PTS | Down | F4 | F4_00596 | *manL* | -2.74 | -2.98 |
| Sugar PTS | Down | F4 | F4_03053 | *-* | -3.01 | -3.15 |
| Sugar PTS | Down | F4 | F4_02811 | *-* | -3.03 | -3.18 |
| Sugar PTS | Down | F4 | F4_03052 | *-* | -3.15 | -3.29 |
| Sugar PTS | Down | F4 | F4_03060 | *-* | -1.59 | -3.31 |
| Sugar PTS | Down | F4 | F4_00260 | *pts5ABC* | -2.94 | -3.35 |

Notes: “-” indicates that the gene name is not assigned.

**Table S5. Differentially expressed genes (DEGs) encoding aminotransferases in strain F4 under ethanol exposure.**

| **Gene_ID** | **Gene Name** | **log_2_FC**  **(24 h)** | **Regulation**  **(24 h)** | **log_2_FC_**  **(48 h)** | **Regulation**  **(48 h)** | **Description** |
| --- | --- | --- | --- | --- | --- | --- |
| F4_01149 | *araT* | 4.70 | Up | 4.35 | Up | Aminotransferase |
| F4_02389 | *patB* | 2.91 | Up | 2.07 | Up | Aminotransferase, class I |
| F4_02136 | *ilvE* | 2.22 | Up | ns | ns | Branched-chain amino acid aminotransferase |
| F4_02085 | *iscS2* | 1.91 | Up | ns | ns | Aminotransferase class V |
| F4_01563 | *aspB* | 1.55 | Up | ns | ns | Aminotransferase |
| F4_00807 | *aspC* | -1.64 | Down | -1.86 | Down | Aminotransferase |
| F4_02517 | *patB* | -2.58 | Down | -2.40 | Down | Aminotransferase, class I |
| F4_02333 | *araT2* | -2.62 | Down | -2.01 | Down | Aminotransferase |
| F4_02788 | - | -3.36 | Down | -2.17 | Down | Aminotransferase, class I |

Notes: ns represents no significant difference; “-” indicates that the gene name is not assigned.

**Figure S1. Colony counts for BNCC 194390 and CICC6009 with versus (vs.) without ethanol stress.** Each sample was analyzed in triplicate.

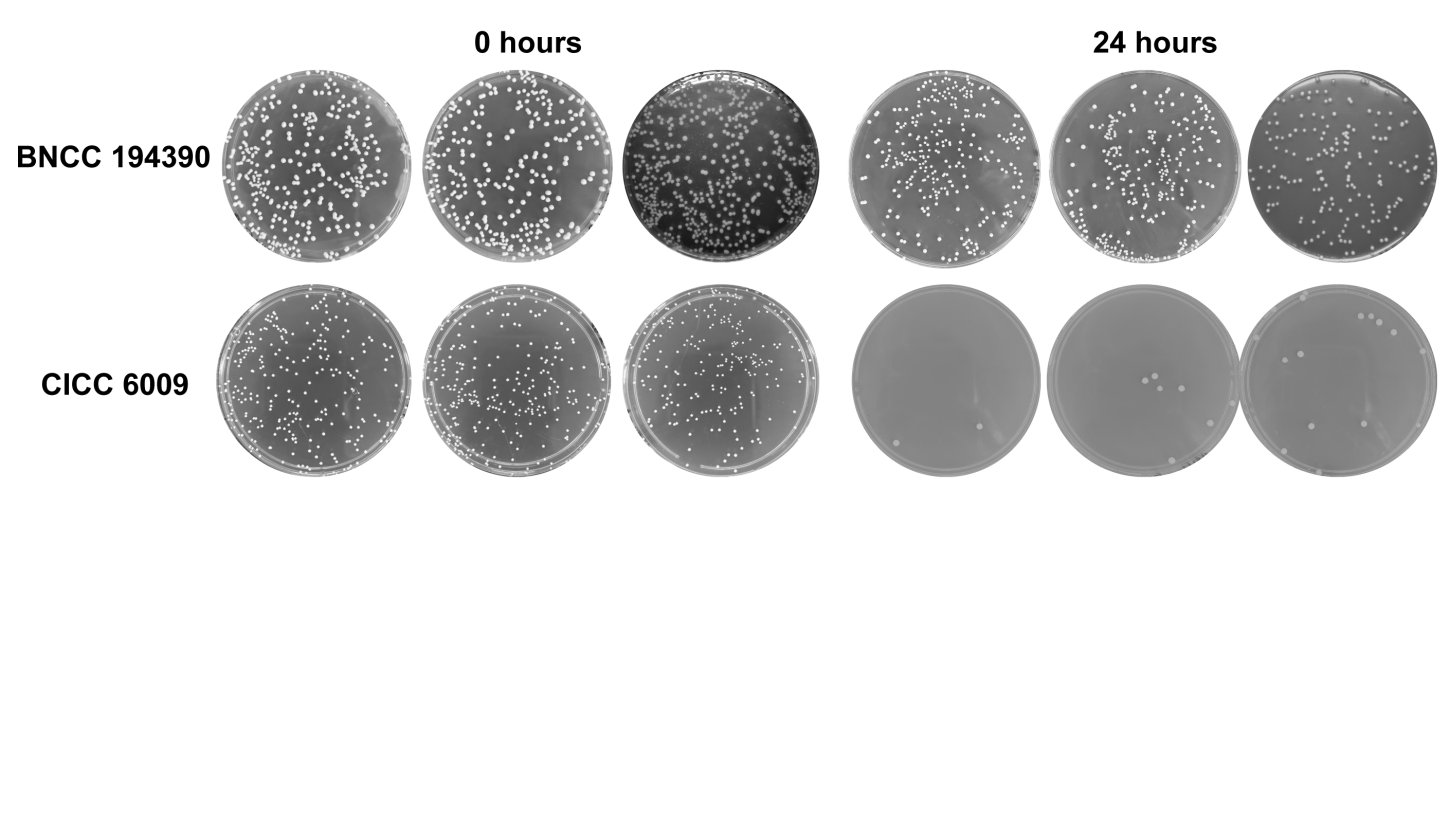

**Figure S2. Metabolic vitality landscape of the pit mud microbiome.** Distribution of single-cell CDR values for over 1,000 microbial cells, with the sorting threshold for the most active 1% indicated.

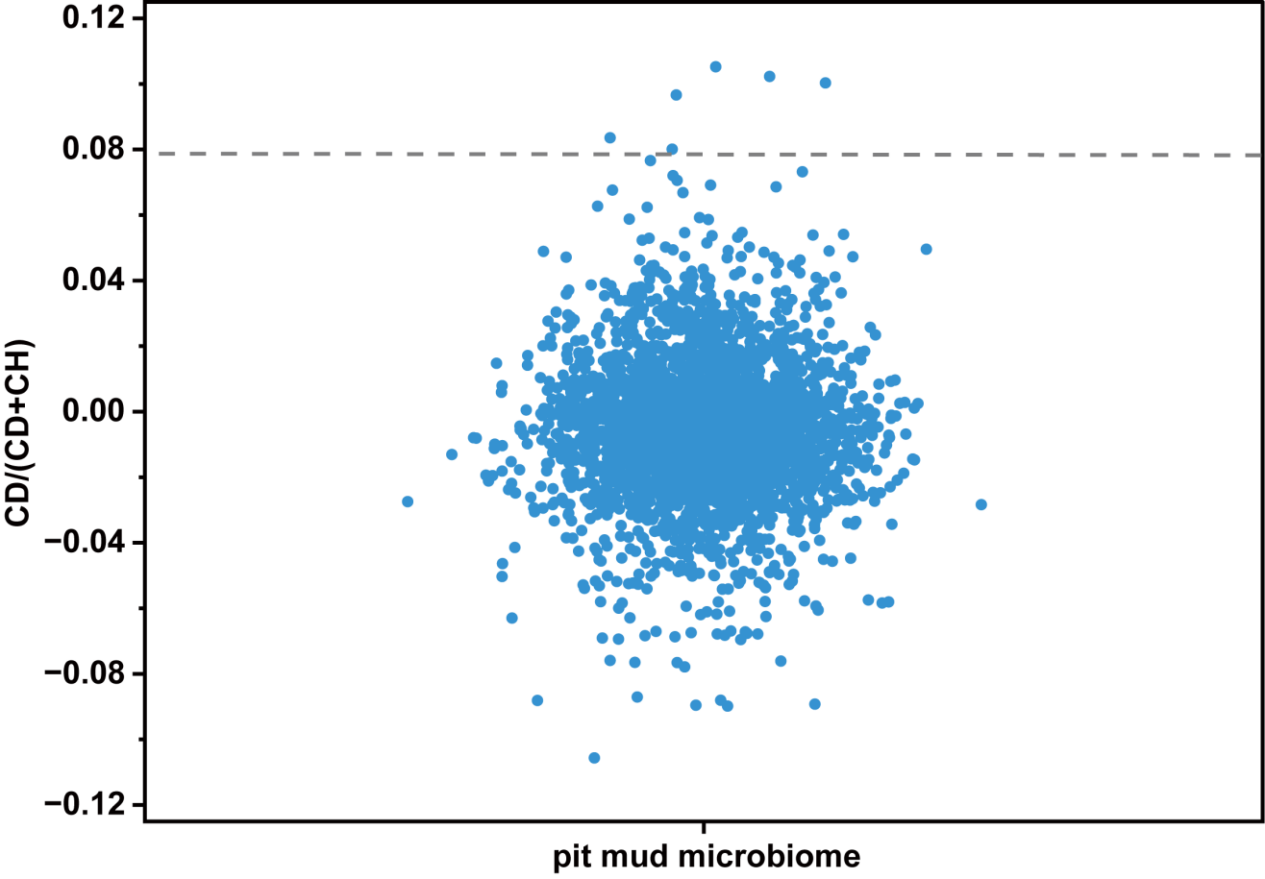

**Video S1. Visualizing fluid stability in the sorting process.** This video shows the stable laminar flow profile within the microfluidic chip during operation, ensuring accurate sorting decisions.

Video S1.mp4

**Video S2. High-efficiency single-cell trapping in action.** This video demonstrates the rapid and precise optical trapping of individual cells by the 1064 nm laser for subsequent sing-cell sort.

Video S2.mp4
